# Representation Methods of Transcriptomics with Applications in Neuroimmune Biology

**DOI:** 10.64898/2026.04.03.716238

**Authors:** Mohammad Abbasi, Santiago Ochoa Zermeño, Mauri D. Spendlove, Zeinab Tashi, Christopher L. Plaisier, Benjamin B. Bartelle

## Abstract

Interpretable representations of gene expression are used to define cellular identities and the molecular programs active within cells, two related, but distinct phenomena. In the case of microglia, a cell type with high transcriptomic, functional, and morphological heterogeneity, the predominant representation of transcriptomic data presumes the adoption of distinct molecular identities, despite a lack of easily separable transcriptional states. Here, we explore alternative transcriptomic representations by comparing two single-cell analysis methods: differential expression analysis for identities and co-expression network analysis for molecular programs. For microglia, co-expression network analysis identifies highly significant functional ontologies not resolved by differential expression analysis. The identified co-expression modules are preserved across transcriptomic datasets and suggest reducible functional programs that activate and modulate depending on context. We conclude that co-expression analysis constitutes a best practice for single cell analysis of an individual cell type and describing microglia function as concurrent molecular programs offers a more parsimonious model of microglia function.

## INTRODUCTION

Microglia at any one moment can phagocytose synapses, secrete both pro- and anti-inflammatory mediators, and remodel their own morphology. In some cases, they do some or all of these things simultaneously, underscoring a remarkable functional plasticity and a spectrum of states that shift depending on intrinsic heterogeneity and microenvironmental cues. Transcriptomic approaches only further elaborate the spectrum of novel microglial states associated with development, aging, and disease (Chhatbar et al., 2026; Hammond et al., 2019; Zheng et al., 2021). This functional complexity has historically complicated efforts to develop consistent nomenclature and classification schemes for microglia.

One approach is to define microglia according to context such as Disease-Associated Microglia (DAM), a transcriptional state first identified in mouse models of neurodegeneration, particularly Alzheimer’s Disease (AD) (Samant et al., 2024). Functionally, DAM are characterized by enhanced phagocytic and lipid metabolism pathways and are found spatially clustered around amyloid-β plaques in AD models. Similar and overlapping reactive states have also been described, including Activated Response Microglia (ARM), the Microglial Neurodegenerative Phenotype (MGnD), and White Matter-Associated Microglia (WAM), which largely share the core transcriptomic signature and dependency on the TREM2-ApoE signaling axis (Sun et al., 2024). However, these functional classifications remain only partially compatible with molecular definitions. Single cell transcriptomics at best reveal enrichment of context dependent microglia in separable clusters, with high heterogeneity (Chhatbar et al., 2026; Hammond et al., 2019; Zheng et al., 2021).

The most common analytical pipeline for scRNA-seq data focuses on the identification and annotation of discrete cell populations. This process typically begins with dimensionality reduction, using techniques such as Principal Component Analysis (PCA) with Uniform Manifold Approximation and Projection (UMAP) to reduce the high-dimensional expression matrix of 3-5k genes down to ∼10 components, for projection onto a 2D plot. At the PCA level, unsupervised clustering algorithms (e.g., graph-based methods like Louvain or Leiden) (Satija et al., 2015) are applied to group cells based on the global similarity of their transcriptomic profiles. When clustering, the central assumption is that cells are differentiated, and conversely that co-location in transcriptional space equates to being of the same type. Differential gene expression analysis (DEA) builds off this assumption to identify genes that are significantly enriched in each cluster relative to the others. Ideally, “marker genes,” serve as a minimal representation of biological identity to each cluster. This cell-centric approach is exceptionally effective for cataloging the cellular composition of differentiated cell types within a tissue and defining the unique transcriptional features of each constituent population (Costa-Silva et al., 2017). While DEA excels at pinpointing gene markers between cell types, it does not capture the underlying structure of coordinated transcriptional programs that support the function of a single cell.

Co-expression network analysis (CNA), does not assume differentiable cells, instead examining how sets of transcripts co-vary across cells or conditions. The core assumption behind CNA is that genes with similar expression profiles across a range of conditions or cell states are likely to be functionally linked. A gene-wise similarity matrix, usually based on pair-wise correlations, can then be conceptualized as a network where nodes represent genes, and the strength of their co-expression determines the weight of the edges connecting them. Within these networks, densely interconnected groups of genes can be identified. These “modules,” are often interpreted to contain collections of genes that are co-regulated or that cooperate to perform specific biological functions, often with validation from gene ontology or transcription factor databases (Fu et al., 2019). When applied to a diverse cell population CNA returns spuriously averaged networks where apparent global correlations are merely artifacts of underlying population structure, termed Simpson’s paradox (Wang et al., 2021). For a single cell type however, CNA is highly effective for generating hypotheses about coordinated gene function and pathway activity not otherwise captured by DEA.

Because of the different underlying assumptions between DEA and CNA, the same cellular heterogeneity that confounds molecular definitions of state is the basis for identifying transcriptional modules. To explore the utility of the two methods, we draw from large publicly available microglia scRNA-seq datasets, including The Allen Brain Cell Atlas (ABCA) and microglia optimized protocols (Chhatbar et al., 2026; Hammond et al., 2019; Jin et al., 2025; Yao et al., 2023). We compare the ontological enrichment of genes by the two methods and characterize clusters vs modules and compare how the two methods differ in the information they offer.

## RESULTS

### Differential expression analysis of Allen Brain Cell Atlas microglia fails to define robust subtypes with functional specialization

From the publicly available scRNA-seq datasets in the ABCA, we selected all microglia isolated using the 10X-v3 technology, which included ∼80,000 cells taxonomically labeled as a single subclass (“334 Microglia NN”). In a 2D UMAP, these microglia appear contiguous, but high dimensional clustering resulted in five distinct transcriptional groups (**Figure 1a**). DEA between groups identified patterns of gene enrichment, however markers were not unique to their cluster (**Figure 1b**). The most significant overlap was between clusters 2 and 3, which aligned with their overlap in UMAP space, and suggested only a minor transcriptional difference.

**Fig 1.**
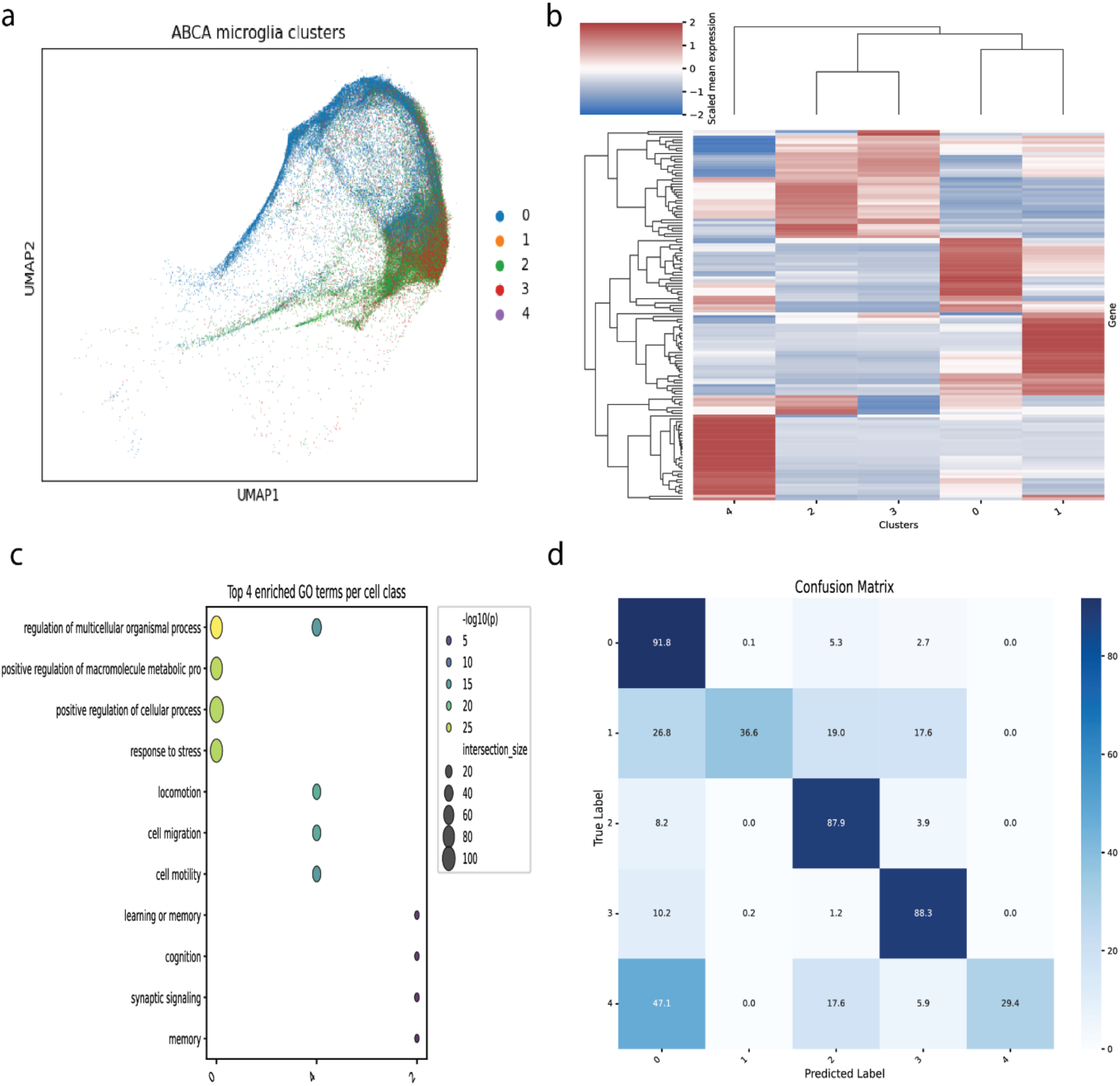
Differential expression analysis of ABCA microglia poorly separable subgroups lacking functional distinction. **(a)** UMAP plot of ∼80,000 microglia cells from the ABCA, colored by five data-driven clusters identified through unsupervised analysis. **(b)** Heatmap of scaled mean expression for the top differentially expressed genes across the five microglial clusters. **(c)** Gene Ontology enrichment analysis of marker genes for each cluster. **(d)** Confusion matrix for a Random Forest classifier trained on cluster DEGs and tested on independent Microglia dataset collected by 10X-V2 technology.

Gene Ontology (GO) enrichment analysis of differentially expressed genes provided minimal functional insights (**Figure 1c**). Clusters offered general terms such as “cell motility” (Cluster 4) and neuron-associated functions (Cluster 2). No significant GO terms were found for Clusters 1 or 3, indicating that their transcriptional signatures do not map cleanly onto well-defined biological processes or are not significantly distinct.

To test the generalizability of DEA we trained a classifier to predict cluster labels against an independent microglia dataset, generated by ABCA with a different technology (10X-v2) (**Figure 1d**). While 3 of 5 clusters were well-predicted (≥87% accuracy), the classifier performed poorly for Cluster 1 (37% accuracy) and Cluster 4 (29% accuracy), with Cluster 4 more likely to be classified as Cluster 0. By this analysis, only one separable cluster had sufficiently distinct expression to suggest biological function and be distinguishable in an independent dataset.

### Co-expression network analysis identifies a repertoire of microglial functions

Analyzing the same dataset with CNA identified five distinct modules of co-expressed genes (**Figure 2a**). Each module represents a discrete transcriptional program, as illustrated by the interconnected network graph of the “MG-M2” module (**Table 1, Figure 2b**). To test the generalizability of identified modules, we reproduced the CNA on the ABCA validation dataset and tested how well the modules were preserved. All modules were statistically reproducible, with four being highly preserved (Z-score > 10) and only one being moderately preserved (MG-M5, Z-score > 2) (**Figure 2c**).

**Table 1.** Modules identified by CNA and their description based on the significant GO terms.

| Module | Compiled GO Name | Hub Genes |
| --- | --- | --- |
| MG-M1 | Classical Inflammation<br>Module | A |
| MG-M2 | Phagocytosis & Antigen<br>Display Module | B |
| MG-M3 | ECM Remodeling Module | C |
| MG-M4 | Interferon Response Module | D |
| MG-M5 | Gliosis Module | E |

**Fig 2.**
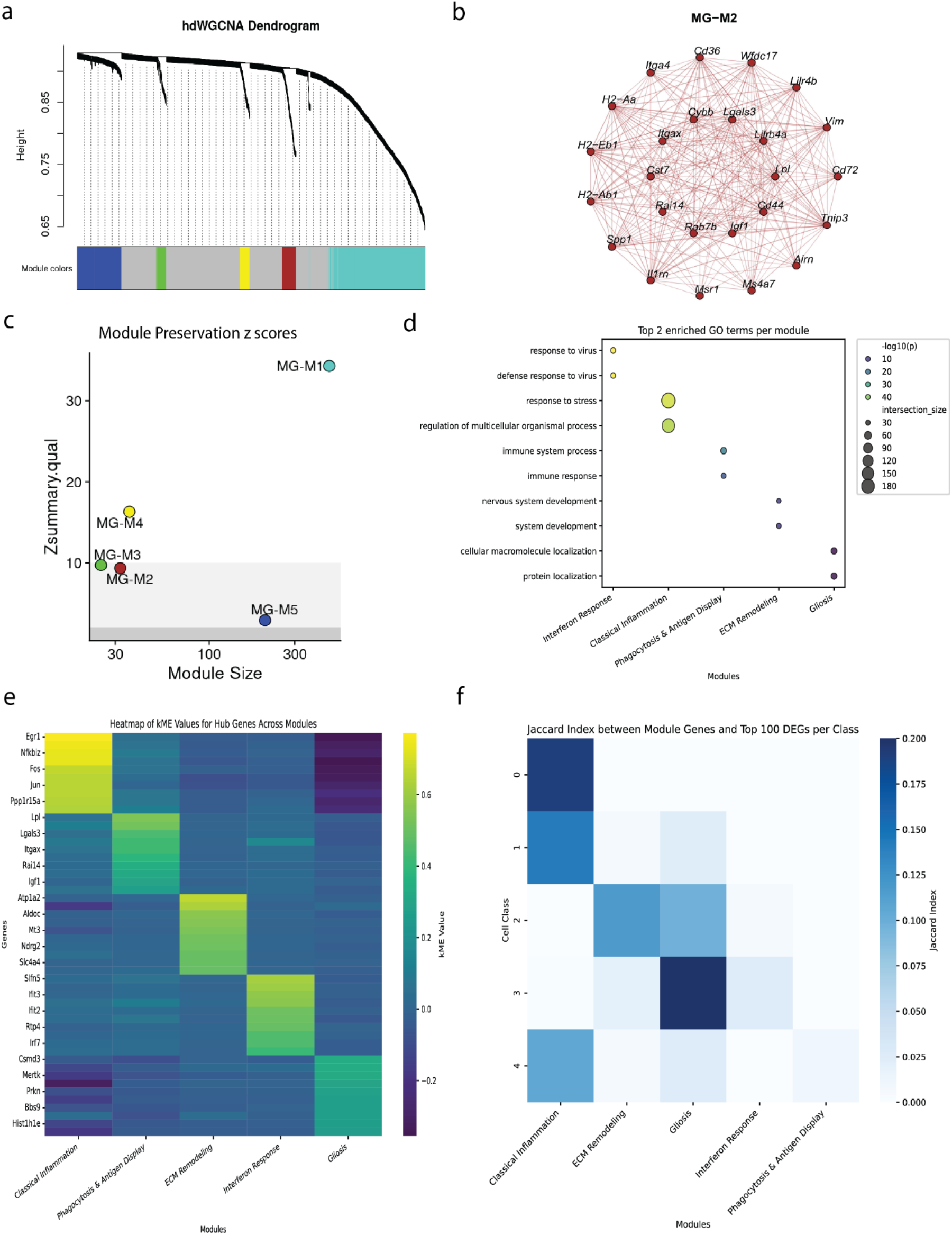
Co-expression network analysis defines robust and functionally coherent microglial programs. **(a)** Hierarchical clustering dendrogram of genes within the microglia population. The color bar indicates the five distinct co-expression modules identified. **(b)** Network plot visualizing the co-expression relationships within an example module (“MG-M2”). **(c)** Module preservation analysis showing Z-summary scores for all five modules in an independent dataset (10X-V2). **(d)** Gene Ontology enrichment analysis for each module. Dot size represents gene count; color indicates statistical significance. **(e)** Heatmap of module connectivity for hub genes across all modules. **(f)** Jaccard index heatmap measuring the fractional overlap between module genes and the top DEGs for each cluster.

To characterize the functions these modules represent, we performed GO enrichment analysis and identified numerous distinct terms for each module. Unlike for DEA, all modules had abundant and specific GO terms. We assigned a descriptive name to each module based on the most significant terms and cross referenced the most correlated genes with the literature (**Figure 2d**). Notably, all modules were associated with cell stress and inflammation, which was expected, given that enzymatic dissociation for scRNA-seq leads to an artifactual transcriptional signature, especially in microglia (Marsh et al., 2022).

We then asked how well each module could be defined using only a subset of the most correlated genes, called the module eigengene (ME) (**Figure 2e**). Each ME, by definition, was highly and specifically correlated with its own module’s activity. Seeing many familiar genes to DEA approaches (lile, lpl, and lgals3) appear as hub genes led us to ask how much overlap there was between MEs and differentially expressed genes (DEGs) (**Figure 2f**). Using a Jaccard index to measure the overlap, some modules clearly correspond to the DEA signals of specific clusters; for example, the “Classical Inflammation” module is largely composed of genes identified as markers for clusters 0, 1, and 4, the same modules that failed to differentiate on a per cell basis using a classifier. Similarly, the “Gliosis” module aligns with markers for Clusters 2 and 3, which had the highest transcriptional overlap. The two modules “Interferon Response” and “Phagocytosis & Antigen Display” showed minimal overlap with the DEGs of any cluster. Modules derived using CNA captured a biological signal that could not be otherwise detected and resolved ambiguous results generated by DEA.

The utility of CNA becomes clear when examining the correlative expression of MEs across previously defined clusters (**Figure 3a**). Here the “Classical Inflammation Module” (MG-M1) shows a clear gradient of activity: it is most highly expressed in cluster 0 and shows lower, but still present, activity in the clusters 1 and 4, but no activity in clusters 2 and 3. This reframes the “confusion” previously seen in the DEA classifier as a reflection of a biological continuum (**Figure 1d**). Most critically, CNA uncovers biological signals that are invisible to DEA. The “ECM Remodeling” (MG-M3) and “Interferon Response” (MG-M4) modules are rarely expressed in all clusters. Since their expression does not vary significantly between the cluster groups, DEA fails to identify them.

**Fig 3.**
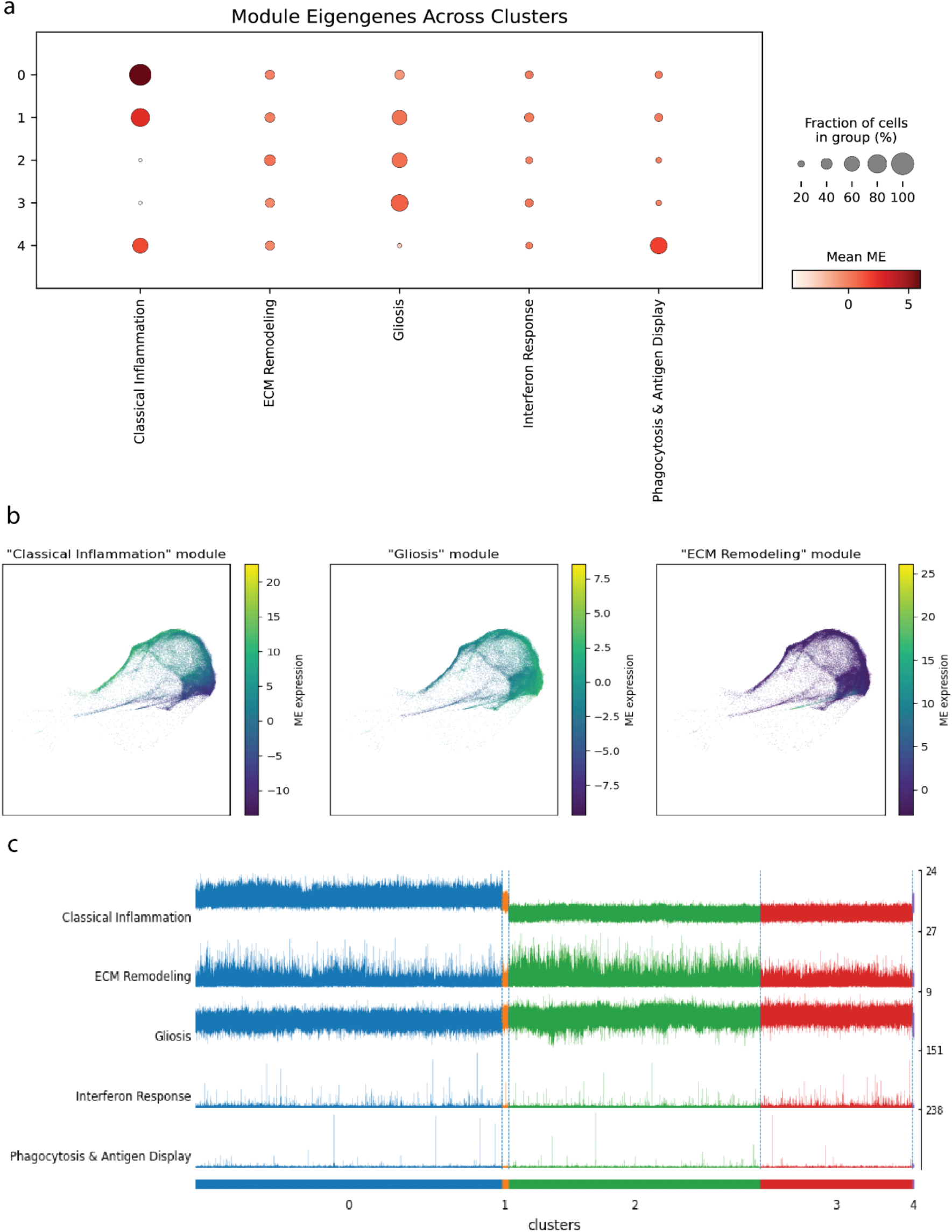
Characterization of microglial modules reveals state-specific and shared functional programs. **(a)** Track plot visualizing the expression of each module eigengene across individual cells, grouped by DEA clusters. **(b)** UMAP projections showing the overlapping and graded expression of three representative MEs. **(c)** Dot plot of the mean module eigengene expression across the five previously defined DEA clusters.

To qualitatively analyze the heterogeneity and overlap of microglial modules, we projected the most abundantly expressed MEs onto the ABCA microglia UMAP embeddings (**Figure 3b**). A final comparison between the five functional modules and clustering at a single-cell level highlights the sparse expression of “Interferon Response” and “Phagocytosis & Antigen Display” MEs that precludes its detection by clustering (**Figure 3c**).

### Differential expression analysis of microglial gene expression in health and disease

While the ABCA dataset provided a broad overview of microglial heterogeneity, the sequenced cells were collected using a neuron optimized protocol that affects microglial transcriptional programs of basal function (Yao et al., 2023). We turned to data generated using techniques optimized for minimal perturbation of microglia (Hammond et al., 2019). In addition to homeostatic adult microglia, we extracted 2 experimental conditions: 7 days post demyelinating lysolecithin (LC) lesion, and a sham injury (saline injection). After correcting for strong batch effects, the UMAP embedding of ∼30,000 cells show a clear structural shift, with the LC-injury cells occupying a distinct region of the transcriptional space, indicating a response to perturbation (**Figure 4a**).

**Fig 4.**
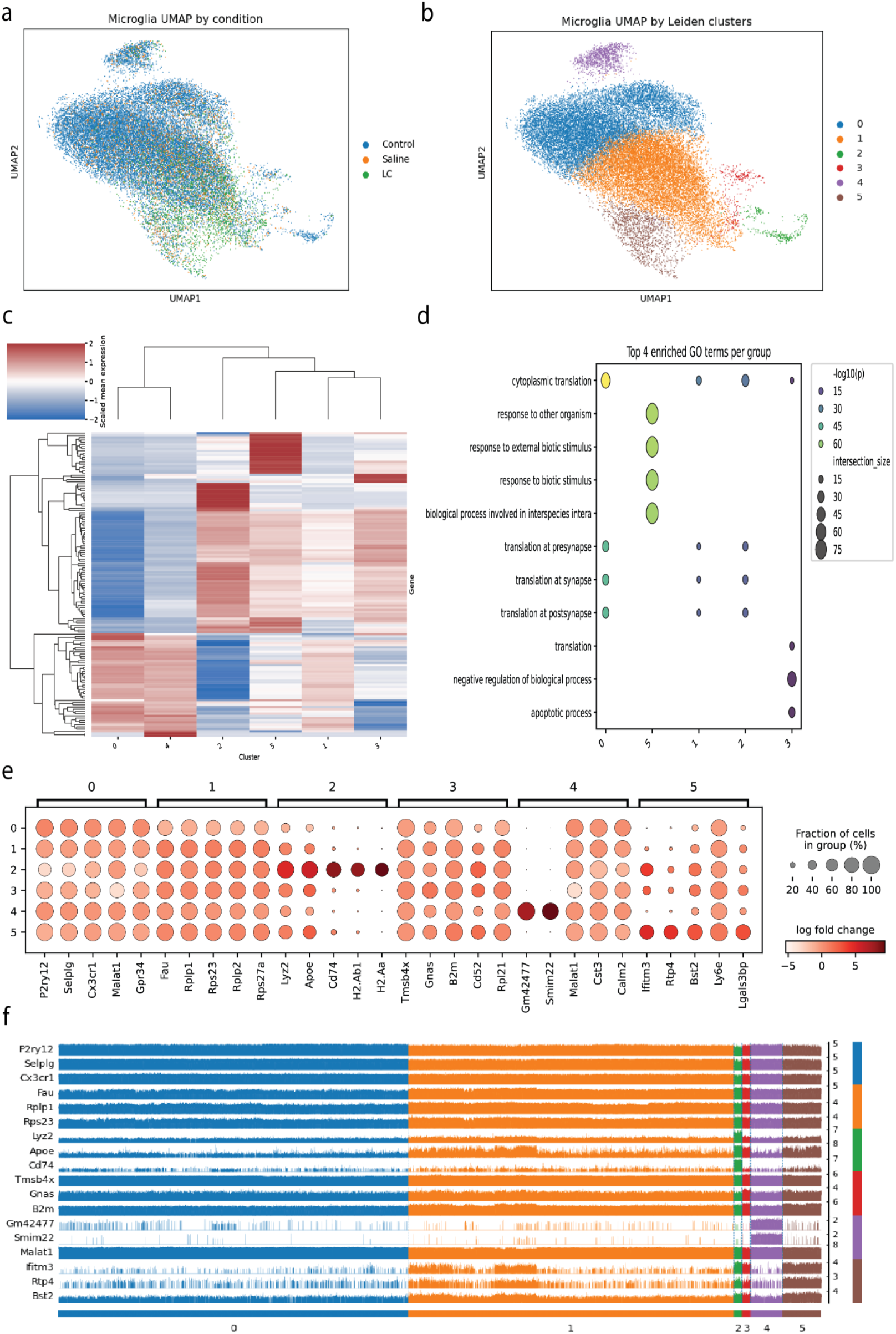
Differential Expression Analysis of Microglia from the Hammond et al. Dataset Reveals a Continuous Activation Spectrum. **(a)** UMAP embedding of microglia from adult mice under control, sham, and lysolecithin (LC) injury conditions. **(b)** The same UMAP colored by six data-driven Leiden clusters. **(c)** Heatmap of scaled mean expression for the top DEGs across the six clusters. (d) Gene Ontology enrichment analysis of marker genes. **(e)** Dot plot of top marker genes expression across clusters. **(f)** Track plot visualizing expression of representative marker genes for different clusters at the single-cell level.

Applying a DEA-based framework produced six clusters with candidate marker genes for each that heavily overlap with previous analysis (**Figure 4b**). However, expression was complex, with blocks of genes showing graded and overlapping expression patterns across the clusters (**Figure 4c**). GO returned significant overlap in the enriched biological processes, particularly between Clusters 0, 1, and 2 (**Figure 4d**). The majority of marker genes were shared across multiple clusters, with only the antigen-presentation genes of Cluster 2 (H2-Aa, H2-Ab1, and Cd74) suggesting a clear function (**Figure 4e**). On a single cell basis, genes identified as markers for one cluster are clearly active in others (**Figure 4f**). For example, Apoe, a top marker for Cluster 2, is also highly expressed in subsets of cells in Clusters 1 and 3. Similarly, Ifitm3, a marker for Cluster 5, is also present in cluster 2. Collectively, these results demonstrate a critical limitation of the DEA framework is identifying distinctive transcriptomic signatures when applied to a plastic cell type undergoing activation.

### Co-expression network analysis of microglial gene expression in health and disease

Applying CNA to Control, Sham (Saline) injury, and LC demyelinating lesion samples independently to avoid normalization steps revealed four modules in control, three in saline, and three in the LC lesion condition (**Figure 5a**). Preservation analysis on random subsets within each condition confirmed that all identified modules were statistically robust and highly preserved within their respective conditions (**Figure 5b**).

**Fig 5.**
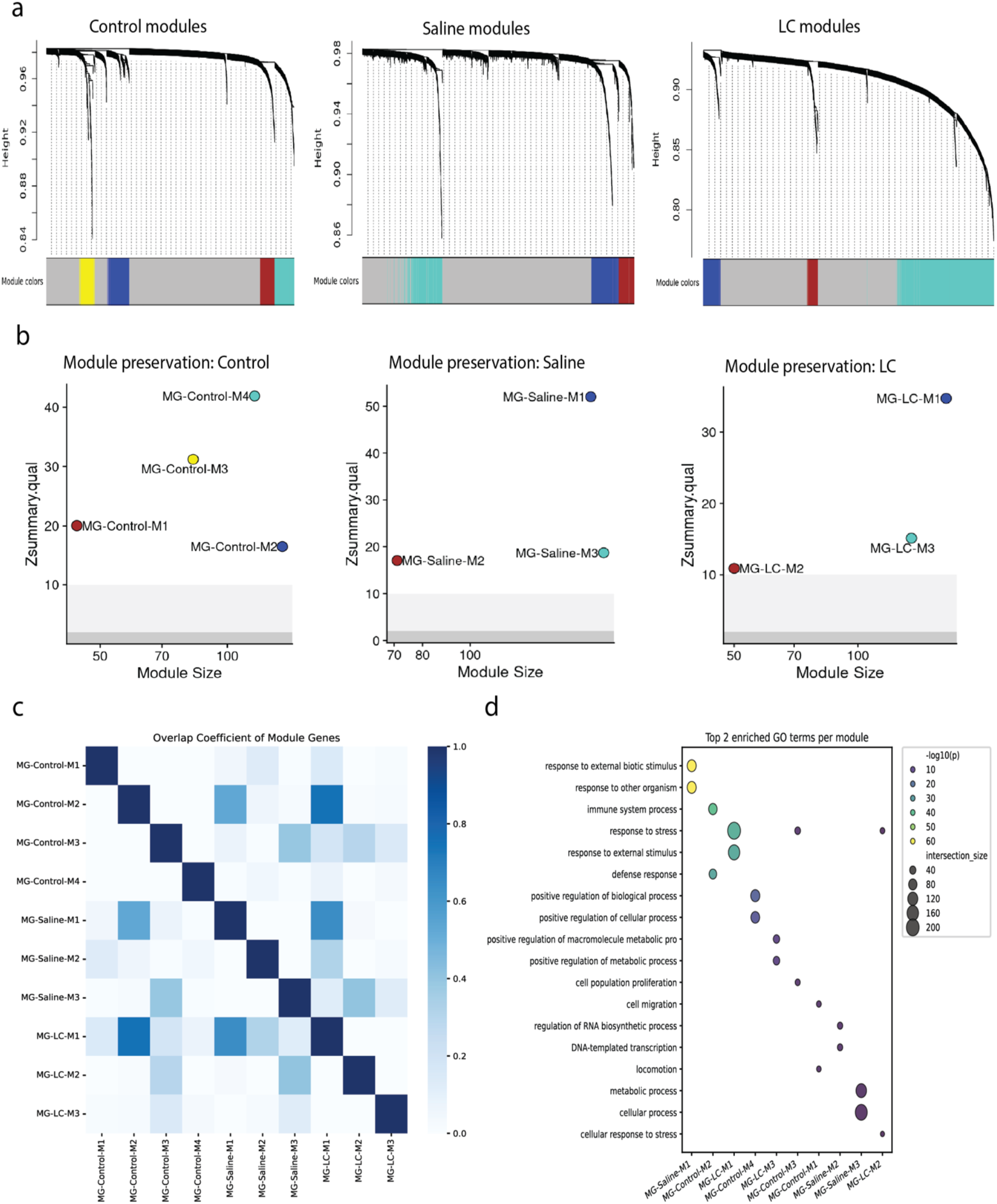
Condition-Specific Network Analysis Reveals Conserved and Specialized Microglial Functional Modules. **(a)** Hierarchical clustering dendrograms showing co-expression modules identified independently in microglia from Control, Saline-injected, and LC-lesioned mice. **(b)** Module preservation analysis confirming the statistical robustness of all identified modules within their respective conditions (all Z-scores > 10). **(c)** Heatmap of the Overlap Coefficient quantifying the gene membership similarity between modules across all three conditions. **(d)** Gene Ontology enrichment analysis for each module, revealing the functional identities of conserved and condition-specific transcriptional programs.

We asked if the modules were repeated, unique to each condition, or reconfigurations of a core set of functions (**Figure 5c**). The genes that made up modules MG-Control-M2, MG-Saline-M1, and MG-LC-M1 highly overlapped, while a second program, MG-Control-M3, MG-Saline-M3 and MG-LC-M2, shared partial overlap between all conditions. Gene Ontology enrichment analysis provided a clear functional identity for these conserved and condition-specific programs (**Figure 5d**). We performed an exhaustive analysis of enriched genes and GO terms to identify distinct transcriptional programs and the microglia biology they represent (**Table 2**)

**Table 2.**
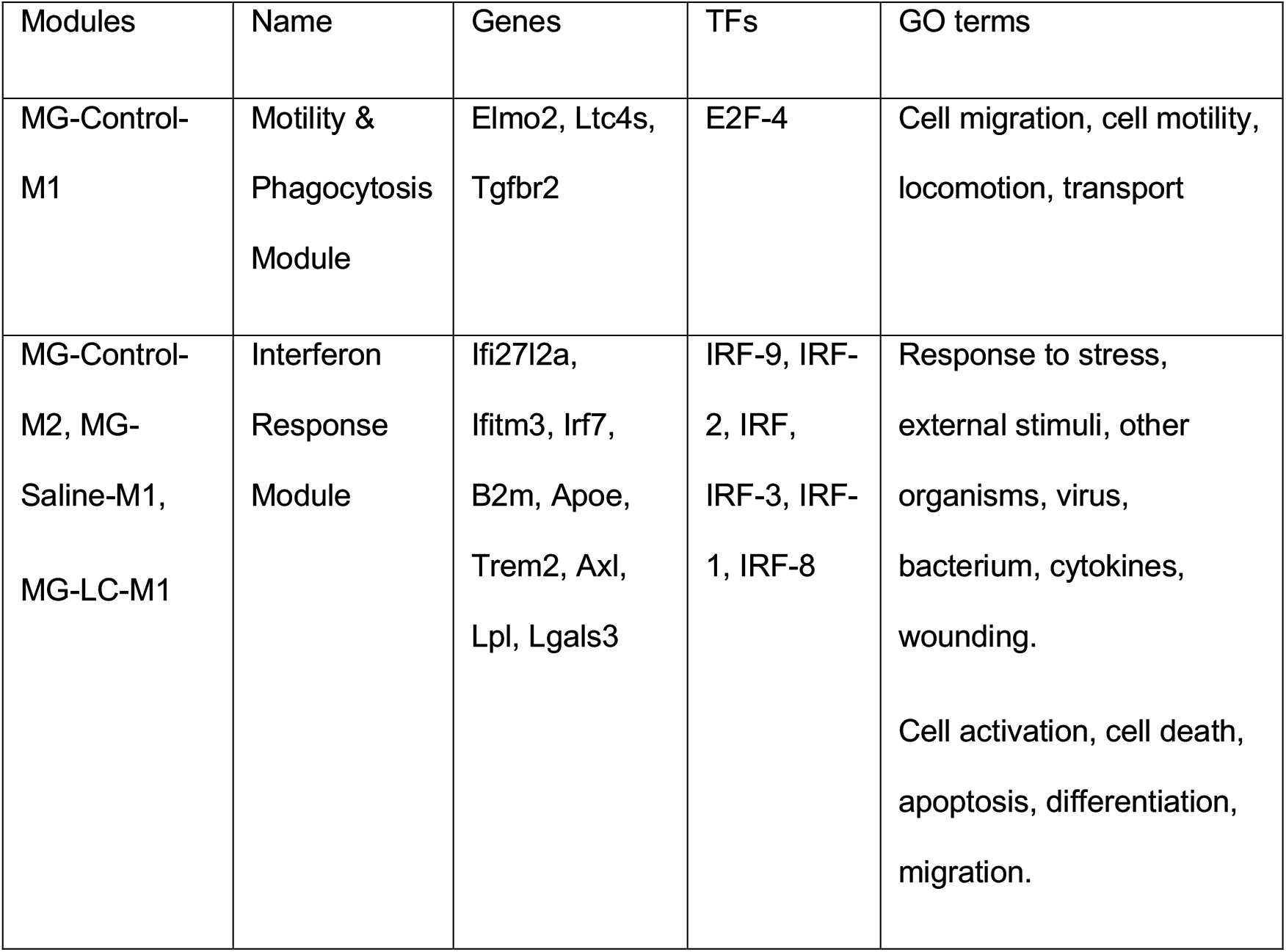
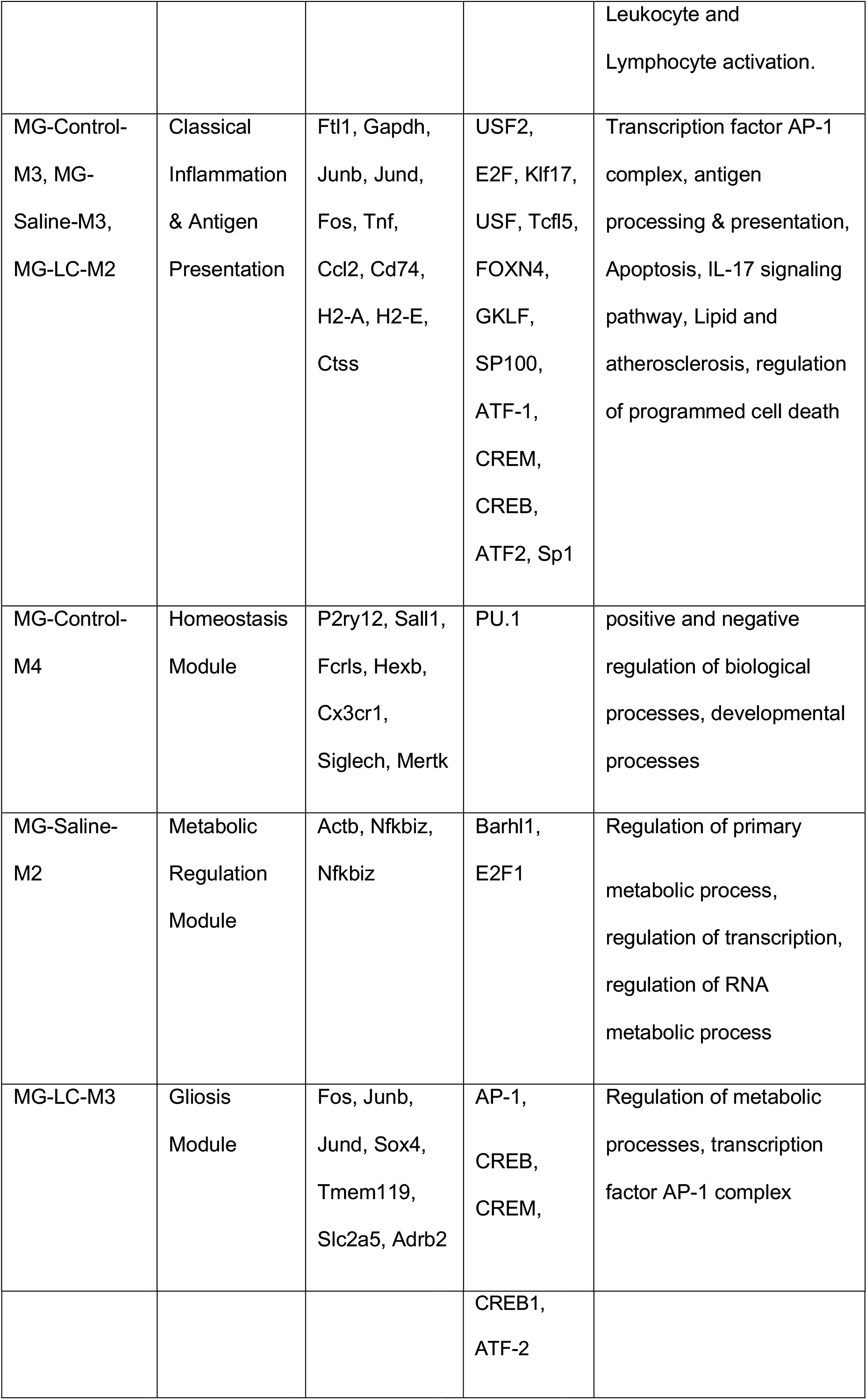
Modules of microglial gene expression with information on hub genes, enriched GO terms and transcription factors.

### A module-based model of microglial transcriptional activation

Consolidating the overlapping modules into a unified set of six core microglial functional programs offers a clear and functionally coherent story of microglial activation (**Figure 6a**).

**Fig 6.**
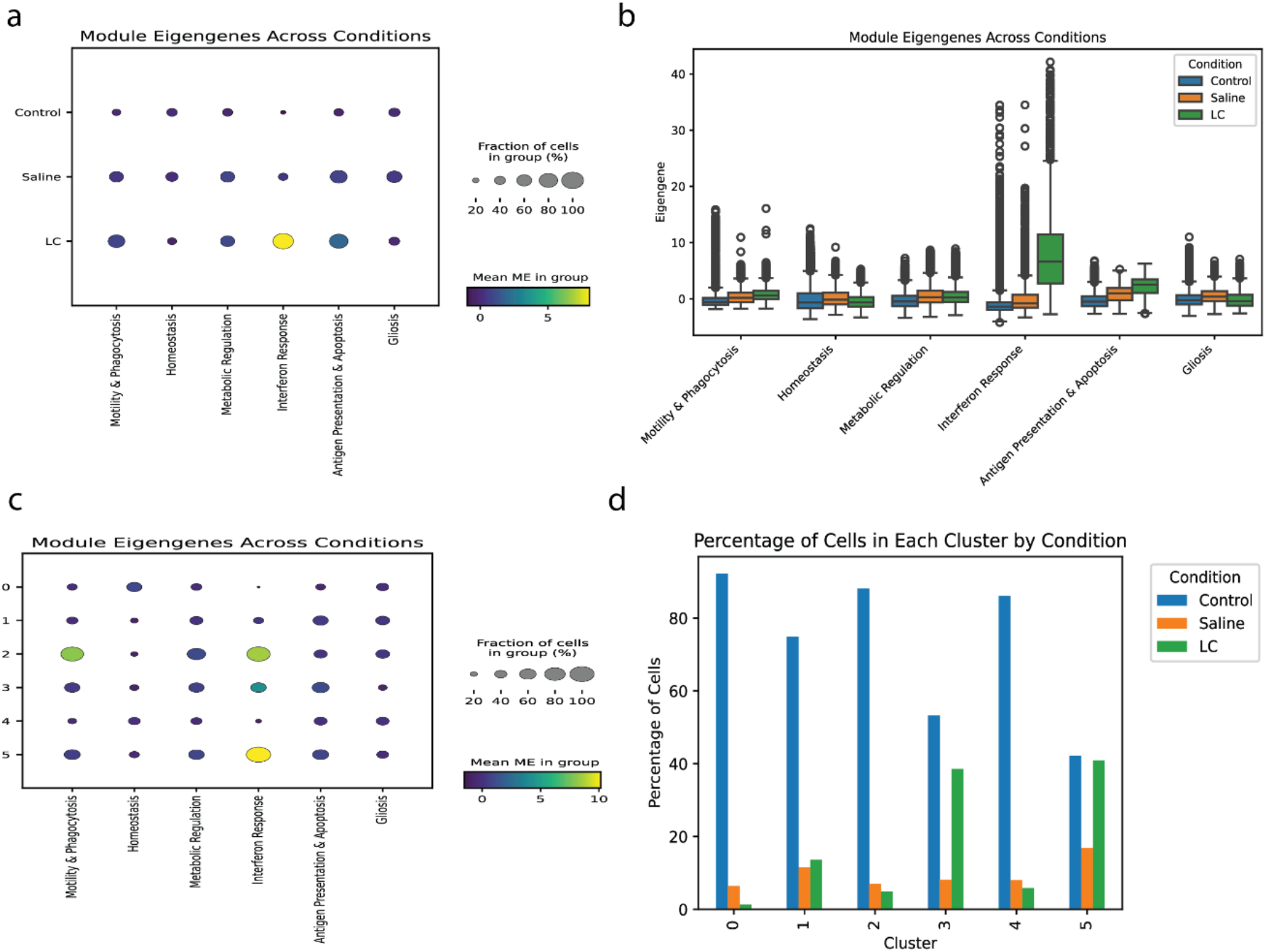
A Module-Based Framework Resolves Microglial Activation States. **(a)** Dot plot showing the mean expression of the six consolidated functional modules across the three experimental conditions. **(b)** Box plot showing the distribution of module eigengene expression across conditions. **(c)** Dot plot showing the mean expression of the six modules across the six previously defined DEA clusters. **(d)** Bar plot showing the percentage of cells from each condition that constitute each DEA cluster.

The “Homeostasis” module, as expected, is most active in control cells and diminished in the saline and LC lesion conditions, tracking the loss of the homeostatic state. Conversely, the “Motility & Phagocytosis” and “Antigen Presentation & Apoptosis” modules show a graded increase in activity from control to saline to the LC lesion. The “Gliosis” module is most prominent in the saline condition. The “Metabolic Regulation” module remains stable across all conditions. Most strikingly, the “Interferon Response” module is most powerfully activated in the LC demyelination condition despite being independently detected across all conditions. While the average expression of the “Interferon Response” module is low in control cells, a small subset of these cells exhibits very high expression (**Figure 6b**). This matches the behavior of the corresponding “Interferon Response” module detected in the ABCA (**Figure 2d**,**3c**).

We sought to reconcile this CNA module-based model with our initial DEA clusters (**Figure 6c**). Modules of Homeostasis, Gliosis, Metabolic Regulation, and Antigen Presentation & Apoptosis are lowly expressed across clusters but from varying degrees of cells. Only the “Motility & Phagocytosis” is uniquely enriched in one cluster and likely contributed to separation.

Projecting the MEs onto the UMAP embedding, the six modules show distinct but highly overlapping patterns (**Figure 7a**). MEs were again heterogeneous on a per cell basis, with the “Homeostasis” module most active in a subset of cells within Clusters 0, 4, and 5. The “Interferon Response” module, a key signature of the LC lesion, is not confined to a single cluster but is strongly expressed by cells in Clusters 2, 3, and 5, and moderately in Cluster 1 (**Figure 7b**).

**Fig 7.**
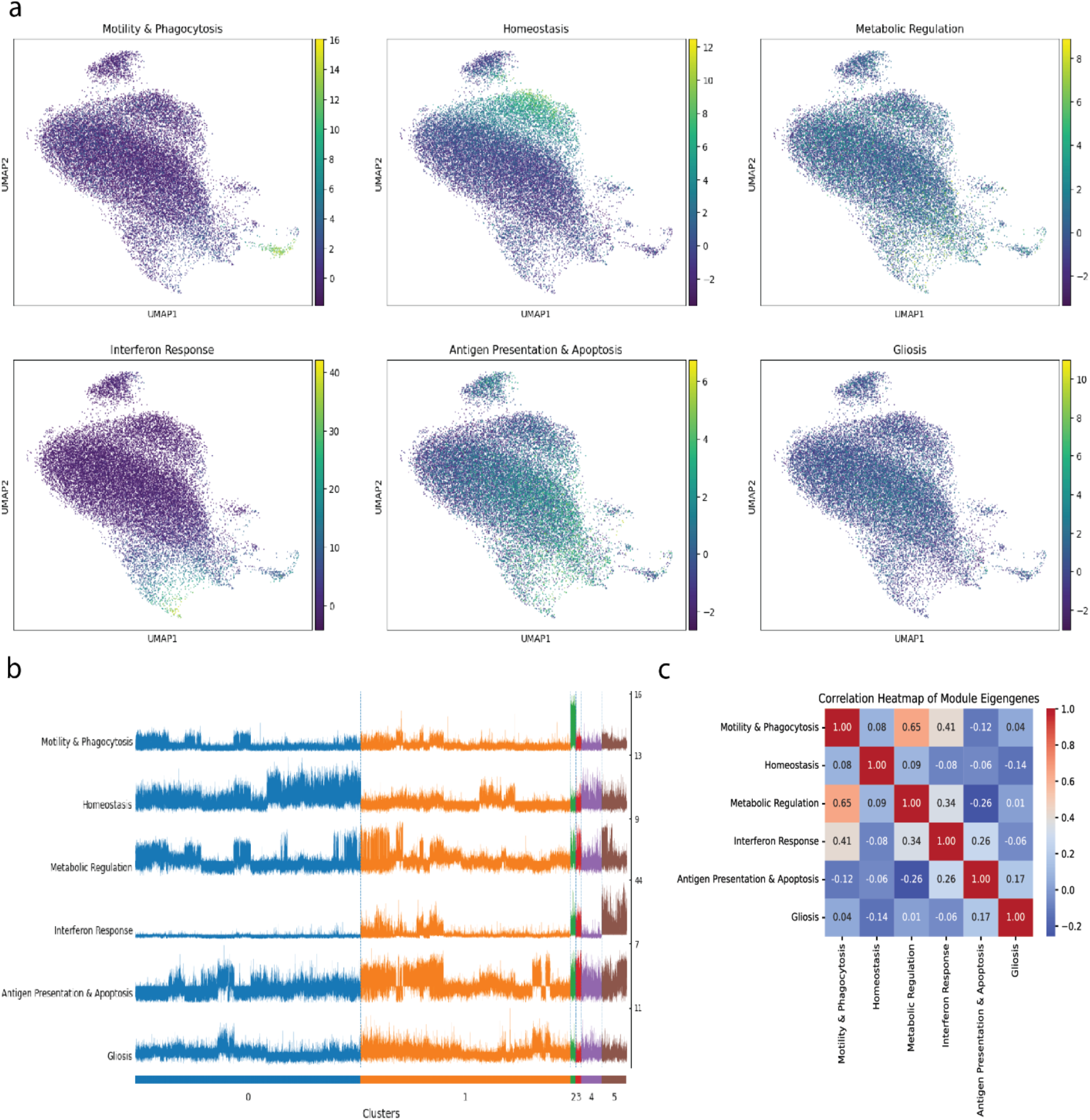
Single-Cell Analysis Reveals Inter-Module Correlations. **(a)** UMAP embeddings of all microglia, with each cell colored by the expression of one of the six consolidated module eigengenes. **(b)** Track plot visualizing the expression of each ME across individual cells, grouped by the six previously defined DEA clusters. **(c)** Heatmap showing the pairwise Pearson correlation between the six module eigengenes across all single cells.

Finally, we investigated whether the activation of these modules occurs independently or in a coordinated fashion by calculating the pairwise correlation between their eigengenes across all cells (**Figure 7c**). This revealed significant relationships between the programs. A strong positive correlation (r = 0.65) was found between the “Motility & Phagocytosis” and “Metabolic Regulation” modules, suggesting a functional link. Conversely “Metabolic Regulation” shared a negative correlation with “Antigen Presentation & Apoptosis.” These inter-module correlations suggest a higher-order regulatory logic governing microglial activation, where the deployment of one functional program can influence the activity of another, a layer of complexity entirely inaccessible through standard DEA.

## DISCUSSION

In this comparison of paradigms for representing transcriptomic information, we have found that CNA can offer more informative representations of biology when describing a single cell type, while DEA best for separating types with stable transcriptional differences. Moreover, the statistical robustness of CNA in our analysis of microglia single cell transcriptomes, supports a model of coordinated and overlapping transcriptional programs. In simplest terms, DEA provides a crucial reductionist view of complex tissues, while CNA offers a systems view, that becomes relevant below the level of cell-type complexity.

The statistical methods underlying DEA do not inherently optimize for expression specificity or uniqueness; its objective is to detect differences between predefined groups, not to ensure exclusivity of expression to one group. In this study, we explicitly tested for specificity, asking whether genes are also uniquely or near-uniquely expressed in target populations, and observed that between clusters, genes are enriched but not exclusive. This behavior is consistent with formal tissue/cell-specificity metrics (e.g., tau score), which quantify a continuum from ubiquitous to highly restricted expression and show that “specificity” is distinct from “differential expression” (Love et al., 2014).

CNA provides a complementary, systems-level view by grouping genes into modules that co-vary across cells. Modules often align with pathways, regulons, or shared functions, even when no single gene is uniquely restricted, reframing subtle subtype differences as combinations of programs rather than as isolated markers. We show empirically that co-expression structure captures functional relatedness across datasets, supporting its role as an organizing principle for fine-scale analysis.

This study includes multiple limitations. First, the analysis is computational and relies on public atlases; conclusions are therefore hypothesis-generating and require orthogonal validation to establish causality. Second, the scope is constrained by adult mouse brain regions represented in current references; generalization to other regions, developmental stages, species, or disease contexts still need to be demonstrated. Third, DEA and CNA are sensitive to upstream choices, normalization, batch handling, clustering/graph construction, which can alter downstream calls; best-practice work emphasizes replicate-aware designs (e.g., pseudobulk for DEA) and robust graph construction before interpretation. Finally, co-expression reflects statistical covariation, not direct regulation; linking modules to mechanism will require multi-omic integration and/or perturbation.

A current limitation of this analysis is that differential expression results were ranked solely by statistical significance, without explicitly incorporating measures of expression prevalence such as the percentage of cells expressing each gene within and outside a cluster. While this ranking strategy ensured consistency across hierarchical comparisons and facilitated direct integration with network-based analyses, it may underrepresent genes that are highly specific to a cell type but expressed in only a small subset of cells. Future iterations of this framework could incorporate multi-criterion ranking schemes that account for both effect size and prevalence, thereby prioritizing genes that are not only significantly differentially expressed but also selectively and consistently detected within their defining populations. Integrating such specificity-aware filtering, alongside metrics like the Tau score, would enhance the biological precision of identified markers and improve correspondence between gene-level and module-level representations of cell identity.

An important consideration in interpreting module activity is the presence of modules or clusters that show little to no detectable expression. Modules with negligible eigengene values in a given cell type likely represent transcriptional programs that are inactive or context-specific, reflecting the modular and selective nature of gene regulation rather than analytical noise. For example, immune- or glia-associated modules would be expected to show minimal activity in neuronal populations, consistent with their restricted biological function. Conversely, clusters that exhibit uniformly low correlations with all modules often correspond to transcriptionally homogeneous expression, where no coherent co-expression structure can be detected at the chosen resolution. Such cases are an inherent aspect of network-based analysis: co-expression modules are designed to capture subsets of coordinated transcriptional variation, not global expression activity. Thus, the absence of eigengene expression is itself informative, delineating the boundaries of program activity across cell types and helping distinguish where specific regulatory processes are active versus silent within the cellular landscape.

A promising future direction emerging from this work is to extend differential expression analysis from individual genes to the modules themselves. Rather than testing each gene independently, one could treat the module eigengene as the unit of analysis and evaluate whether these module-level signals differ significantly across conditions, cell types, or perturbations. This differential module expression approach would directly assess whether coordinated transcriptional programs, rather than isolated genes, are up- or downregulated between biological contexts. Such analyses could reveal global shifts in pathway activity even when individual genes show modest or inconsistent expression changes, thereby improving sensitivity to subtle but functionally coherent alterations. Combining module-level differential testing with traditional gene-level results would provide a hierarchical framework for interpreting transcriptional regulation, linking discrete marker changes to broader pathway reprogramming, and could guide experimental validation toward the most biologically integrated signals of cell-type identity and state transitions.

## METHODS

### Computational analysis workflow

scRNA-seq analysis typically begins with data acquisition, where high-throughput sequencing generates raw reads, and molecular barcodes (UMIs - Unique Molecular Identifiers) are used to tag transcripts originating from individual cells and molecules. Next, data is processed by demultiplexing these barcodes, aligning reads to a reference genome or transcriptome, and quantifying gene expression to produce a count matrix (**Figure 9**). In this count matrix, rows represent genes and columns represent cells (Traversa & Chiara, 2025). Subsequent steps focus on refining this raw data. Quality Control (QC) and filtering are crucial for removing low-quality cells (e.g., those with few detected genes or high mitochondrial gene content, indicative of stress or apoptosis) and uninformative genes. Normalization then adjusts for differences in library size (total number of UMIs per cell) and other technical confounders to make expression levels comparable across cells. Feature selection aims to identify highly variable genes (HVGs) across the cell population, which are presumed to carry the most biological signal and are used for downstream dimensionality reduction.

**Fig 9.**
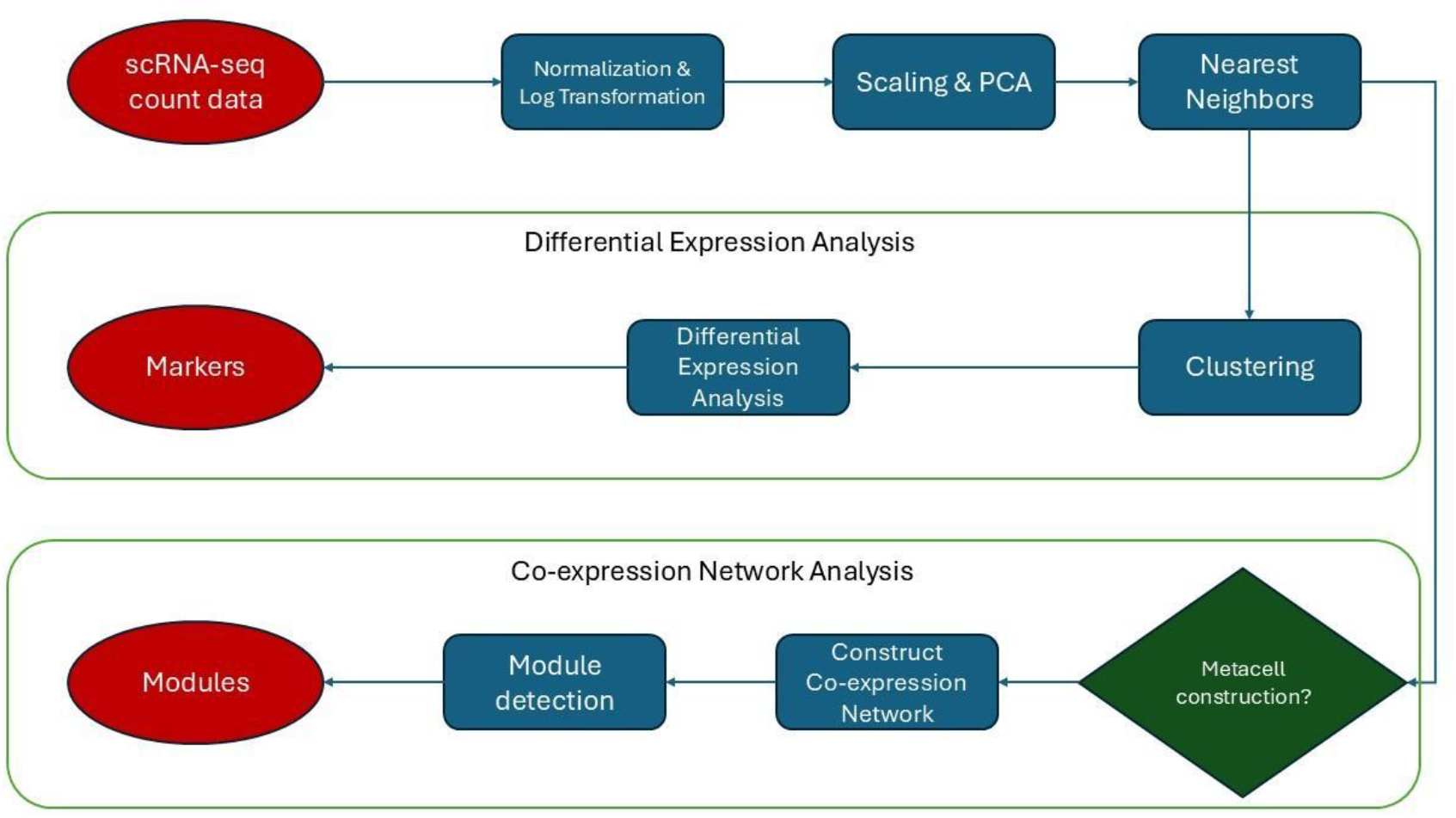
Workflow diagram of scRNA-seq processing steps.

To manage the high dimensionality of the data and reduce noise, Principal Component Analysis (PCA) is applied to the selected HVGs. For visualization and further analysis, non-linear dimensionality reduction methods such as t-distributed Stochastic Neighbor Embedding (t-SNE) or Uniform Manifold Approximation and Projection (UMAP) are commonly used to project cells into a lower-dimensional space (Liu et al., 2026). Based on these embeddings, clustering algorithms (e.g., graph-based methods like Louvain or Leiden applied to a K-Nearest Neighbor graph of cells) group cells with similar transcriptomic profiles. Finally, cell type annotation assigns biological identities to these clusters, often by examining the expression of known marker genes or by using automated annotation tools that leverage reference datasets(Traversa & Chiara, 2025).

For DEA, a statistical model is fitted to the data for each gene; for scRNA-seq data, models based on the negative binomial distribution are commonly used to account for the count nature and overdispersion of the data. This model is then used to perform statistical tests to determine whether the observed differences in gene expression between conditions are statistically significant. Given that thousands of genes are tested simultaneously, a crucial step is the correction for multiple testing (e.g., using methods like Benjamini-Hochberg to control the False Discovery Rate, FDR), to reduce the likelihood of false positives. Finally, the resulting lists of differentially expressed genes are interpreted, often through downstream analyses like gene set enrichment or pathway analysis, to uncover the underlying biological implications (Subramanian et al., 2005).

The construction of a co-expression network typically begins with an expression matrix. First, a similarity measure, most commonly the Pearson correlation coefficient, is calculated for all pairs of genes across the samples to quantify the strength of their co-expression. This results in a large correlation matrix. To emphasize strong correlations and reduce noise from weak or spurious correlations, this matrix is often transformed into an adjacency matrix, typically by applying a power function (soft thresholding) or a hard threshold, where connections (edges) between genes (nodes) are only kept if their co-expression strength exceeds a certain value. This adjacency matrix mathematically defines the network. From this network, clustering algorithms, such as hierarchical clustering, identify modules of highly interconnected genes (Langfelder & Horvath, 2008).

Following is an overview of the main steps of the workflow:

1 Preprocessing scRNA-seq dataset 1a Normalizing counts

1b Log-transforming counts

1c Selecting highly variable genes

1d Scaling

1e Dimensionality reduction

1f Batch correction

1g Clustering

1h Visualization: UMAP

2 Differential Expression Analysis

2a Visualization: MA plot

2b Visualization: Dot plot

2c Visualization: UMAP

2d Visualization: Track plot

2e Visualization: Heatmap of mean expression

2f Visualization: Heatmap of Jaccard Index

2g Specificity: Tau score

2h Coherence: Heatmap of correlations

2i Validation: Classifier model with independent dataset

2j Validation: Biological relevance via gene ontology enrichment analysis

3 Co-expression Network Analysis (CNA)

3a Constructing metacells

3b Selecting soft-power threshold

3c Constructing co-expression network

3d Detecting modules

3e Computing module eigengenes (ME)

3f Computing module connectivity (kME)

3g Visualization: Dendrogram

3h Visualization: Module graph

3i Visualization: Module eigengene expression on UMAP

3j Visualization: Module eigengene expression dot plot 3k Visualization: Module eigengene expression track plot 3l Visualization: Heatmap of Jaccard Index

3m Visualization: Heatmap of connectivity

3n Specificity: Tau score

3o Validation: Statistical robustness via module preservation

3p Validation: Biological relevance via gene ontology enrichment analysis

### 1 Preprocessing scRNA-seq dataset

To ensure high-quality downstream analysis, raw single-cell RNA-sequencing data are processed using Scanpy (Wolf et al., 2018), a scalable and widely used toolkit for analyzing large-scale transcriptomic data in Python.

#### 1a Normalizing counts

The raw UMI count matrix, denoted as (*X* ∈ *R*^*n*×*p*^) where *n* is the number of cells and *p* is the number of genes, contains the number of transcripts (unique molecular identifiers, UMIs) detected for each gene in each cell. Because sequencing depth and capture efficiency vary across cells, the total number of counts per cell can differ substantially, introducing technical biases in the observed expression levels. Normalization is therefore required to make expression values comparable across cells and to ensure that observed differences reflect biological variation rather than sequencing depth.

In this workflow, normalization is performed using library-size scaling, implemented in *Scanpy* as ‘scanpy.pp.normalize_total()’. For each cell *i*, the total number of detected transcripts is computed as:

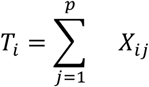

Each gene count *X*_*ij*_ is then divided by this total and multiplied by a constant target value *s* (commonly *s* = 10,000) to obtain normalized counts:

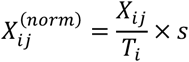

where:

- *X*_*ij*_ = raw UMI count for gene *j* in cell *i*
- *T*_*i*_ = total UMI count for cell *i*
- *s* = target total counts per cell (e.g., 10,000)

This step converts absolute transcript counts into relative abundances, equivalent to transcripts per ten thousand (TP10K). It effectively corrects for sequencing-depth differences while maintaining the internal distribution of gene expression within each cell.

#### 1b Log-transforming counts

After normalization, the count matrix is log-transformed using the natural logarithm function implemented in *Scanpy* as ‘scanpy.pp.log1p()’. The purpose of the log transformation is to stabilize the variance of gene expression values, compress the dynamic range, and reduce the dominance of highly expressed genes.

The normalized expression matrix is denoted as *X*^(*norm*)^ ∈ *R*^*n*×*p*^, where *n* represents the number of cells and *p* the number of genes. For each cell *i* and gene *j*, the log-transformed value is

computed as:

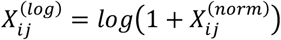

This formulation uses the function *log*(1 + *x*) instead of *log*(*x*) to handle zero counts gracefully and avoid undefined values for genes not detected in a cell. The “+1” acts as a pseudocount, ensuring that every expression value is positive before transformation.

#### 1c Selecting highly variable genes

Many genes exhibit low or near-constant expression across all cells, often representing housekeeping or uninformative transcripts. To reduce dimensionality and focus the analysis on genes that drive biological heterogeneity, highly variable genes are identified using the function ‘scanpy.pp.highly_variable_genes()’.

The principle behind HVG selection is to model the relationship between a gene’s mean expression and its variance across cells. Genes with higher variance than expected for their average expression level are likely to capture meaningful biological signal (e.g., cell-type-specific or state-specific activity), whereas those that follow or fall below the expected variance trend primarily reflect uninformative transcripts or technical noise.

For each gene *j*, we compute the mean and variance across all cells as:

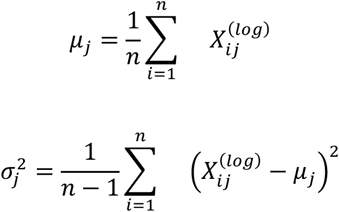

where:

- *n* = total number of cells
- *μ*_*j*_ and 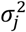 = sample mean and variance for gene *j*

#### 1d Scaling

After selecting highly variable genes, each gene’s expression values are standardized (or z-scored) across all cells so that all genes contribute equally to downstream analyses. This step is performed using ‘scanpy.pp.scale()’. Scaling ensures that each gene has a mean of zero and a variance of one, which is essential for dimensionality reduction algorithms such as Principal Component Analysis that assume features are on comparable scales.

For each gene *j*, the mean and standard deviation of its expression across all *n* cells are calculated as mentioned in section **1c**. Each expression value is then standardized by subtracting the mean and dividing by the standard deviation:

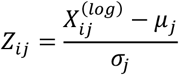

The resulting matrix *Z* contains z-scored expression values, where each gene now has:

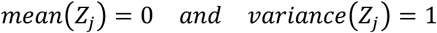

#### 1e Dimensionality reduction

After scaling, the standardized expression matrix *Z* ∈ *R*^*n*×*p*^(where *n* is the number of cells and *p* the number of highly variable genes) is subjected to PCA using ‘scanpy.pp.pca()’. The first step of PCA is to compute the covariance matrix of the scaled data:

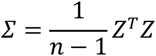

Here, *Z*^*T*^ denotes the transpose of *Z*, and Z ∈ *R*^*p*×*p*^ summarizes how genes co-vary across cells. PCA then performs eigen-decomposition of the covariance matrix:

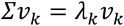

where:

- *v*_*k*_= the eigenvector corresponding to the *k*^*th*^principal component
- λ_*k*_= the eigenvalue indicating the amount of variance explained by that component

The eigenvectors form the **principal component directions**, and the eigenvalues quantify their relative importance.

The original high-dimensional data can then be projected into this reduced-dimensional space as:

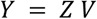

where:

- *V* = [*v*_1_, *v*_2_, …, *v*_*d*_] is the matrix of the top *d*eigenvectors (principal component loadings)
- *Y* ∈ *R*^*n*×*d*^ is the reduced-dimensional representation (typically *d* = 30–50)

Each row of *Y* represents a cell embedded in principal component space, capturing the dominant patterns of transcriptional variation while minimizing noise from lower-variance dimensions.

By construction, the components are orthogonal (uncorrelated), and the total variance explained by the top *d* components is:

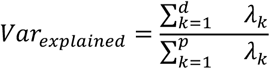

#### 1f Batch correction

To mitigate technical variation due to batch effects (e.g., sequencing runs), Harmony integration (scanpy.external.pp.harmony_integrate) is applied on the PCA-reduced data. Harmony corrects batch differences while preserving the intrinsic biological signal.

#### 1g Clustering

Cell clustering is performed using the Leiden algorithm (scanpy.tl.leiden) based on a k-nearest neighbor graph (scanpy.pp.neighbors). This unsupervised graph-based method identifies groups of transcriptionally similar cells.

#### 1h Visualization: UMAP

To visualize the global structure of the dataset, UMAP is computed using ‘scanpy.tl.umap’. UMAP embeds high-dimensional data into 2D space, revealing the spatial relationship between clusters and aiding in the interpretation of cellular identities.

### 2 Differential Expression Analysis

DEA is done using Scanpy’s ‘tl.rank_genes_groups’ function. It first separates the cells into groups based on given clustering information (e.g., from the Leiden algorithm). Each group is compared against all other cells (or a specific reference group, if specified). For each gene, it checks whether the expression values are consistently higher (or lower) in the group of interest compared to the reference group. It does this one gene at a time, so every gene gets tested separately. Using the Wilcoxon method, the tool ranks the expression values rather than relying on the actual expression numbers. This makes it robust to outliers and well-suited for single-cell data, which can be sparse and non-normally distributed. After testing all genes, the function assigns a statistical score to each gene (like a p-value or test statistic) that reflects how well it distinguishes the target group from others. Genes are then ranked from most to least differentially expressed. In addition to statistical significance, the function also reports log fold change (how much higher the expression is in one group) and adjusted p-values to account for multiple testing (e.g., FDR).

#### 2a Visualization: MA plot

To summarize and interpret differential expression analysis results, MA plots are generated to visualize the relationship between the average expression of each gene and its log fold change between groups. This representation highlights genes that show both strong differential regulation and robust expression levels, allowing rapid identification of potential marker genes.

For each gene *j*, the average expression across the two groups being compared is computed as:

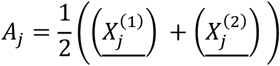

where:

- 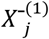= mean normalized expression of gene *j*in group 1
- 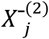= mean normalized expression of gene *j*in group 2
- *A*_j_= average (mean) log expression level for gene *j*

The log fold change (LFC) quantifies the relative difference in expression between the two groups:

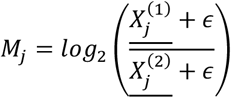

where *ϵ* is a small pseudo-count (e.g., 10^‒6^) added to avoid division by zero.

- Positive *M*_*j*_ values indicate genes upregulated in group 1 relative to group 2.
- Negative *M*_*j*_ values indicate genes downregulated in group 1.

The MA plot displays each gene as a point with coordinates (*A*_*j*_, *M*_*j*_), where the **x-axis** represents average expression (*A*_*j*_) and the y-axis represents log fold change (*M*_*j*_).

Genes that deviate strongly from *M*_*j*_ = 0 correspond to those with substantial differences in expression between groups.

#### 2b Visualization: Dot plot

To summarize and compare the expression of top-ranked differentially expressed genes across multiple clusters or cell types, dot plots are generated using the function ‘scanpy.pl.rank_genes_groups_dotplot’. This visualization integrates both expression magnitude and detection frequency, providing a compact and interpretable overview of gene-expression patterns across groups.

In a dot plot, each row corresponds to a gene and each **column** to a cluster (or cell type). Two main quantities are represented simultaneously:

1. Dot size: the fraction of cells in a cluster that express the gene above a defined threshold (i.e., detection rate).
2. Dot color: the average expression level of that gene within the cluster, typically scaled or transformed (e.g., log_10_ or z-score) to allow comparison across genes.

Formally, for a gene *j* and a cluster *c*:

- The detection fraction (dot size) is computed as

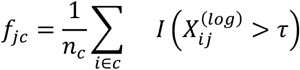

where:

- *n*_*c*_= number of cells in cluster *c*
- *I*(·)= indicator function (1 if true, 0 otherwise)
- *τ*= expression threshold (often 0)
- *f*_*jc*_= proportion of cells in cluster *c*expressing gene *j*
- The average expression (dot color) is computed as

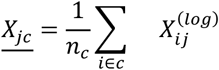

where 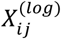 is the log-transformed expression of gene *j* in cell *i*.

The resulting plot uses color intensity to represent 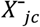 and dot size to represent *f*_*jc*_.

This dual encoding enables simultaneous assessment of how strongly a gene is expressed and how widely it is expressed within each cluster.

#### 2c Visualization: UMAP

Expression of DEGs is visualized on the UMAP embedding to assess spatial restriction and concordance with cluster boundaries using ‘scanpy.pl.umap’. This visualization helps verify whether DEGs are cluster-specific and biologically interpretable.

#### 2d Visualization: Track plot

Track plots are generated using ‘scanpy.pl.rank_genes_groups_tracksplot’ to display the expression of selected marker genes across individual cells of different clusters.

#### 2e Visualization: Heatmap of mean expression

Heatmap of top DEGs across clusters are generated to display the expression of selected marker genes across cells of different clusters.

#### 2f Visualization: Heatmap of Jaccard Index

The Jaccard Index provides a simple yet powerful measure of set similarity, indicating how many DEGs are shared between two clusters relative to the total number of unique DEGs in their union. For any two clusters *a*and *b*, let *G*_*a*_ and *G*_*b*_ denote the sets of DEGs identified for each cluster.

The Jaccard Index between clusters *a* and *b* is defined as:

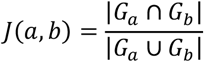

where:

- ∣ *G*_*a*_ ∩ *G*_*b*_ ∣ = number of DEGs shared between clusters *a*and *b*
- ∣ *G*_*a*_ ∪ *G*_*b*_ ∣ = total number of unique DEGs across both clusters
- *J*(*a, b*) ∈ [0,1], with *J*(*a, b*) = 1 indicating complete overlap (identical DEG sets) and *J*(*a, b*) = 0 indicating no overlap

The pairwise Jaccard indices for all cluster combinations are computed to form a symmetric Jaccard similarity matrix:

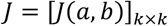

where *k* is the total number of clusters. This matrix is visualized as a heatmap, where the color intensity encodes the value of *J*(*a, b*).

#### 2g Specificity: Tau score

To quantitatively assess how specifically each gene is expressed across clusters, the **Tau score** is computed. This metric measures the degree to which a gene’s expression is concentrated in a single cluster versus being distributed broadly across many clusters. It provides a continuous measure of expression specificity, ranging from 0 (ubiquitous expression) to 1 (perfectly specific expression).

For a given gene, let *x*_*i*_ denote its average expression in cluster *i, x*_*max*_ the maximum expression value across all clusters, and *n* the total number of clusters.

The Tau score is defined as:

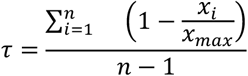

where:

- *x*_*i*_ = average expression of the gene in cluster *i*
- *x*_*max*_ = *max* (*x*_1_, *x*_2_, …, *x*_*n*_) = maximum average expression across all clusters
- *n* = number of clusters

The Tau score is bounded between 0 and 1:

- *τ* = 0: the gene is uniformly expressed across all clusters (no specificity).
- *τ* = 1: the gene is perfectly specific, expressed in only one cluster.
- Intermediate values indicate partial specificity: genes that are enriched but not exclusive to one cluster.

#### 2h Coherence: Heatmap of correlations

Heatmaps are generated to display pairwise correlations among top differentially expressed genes within each cluster. This visualization highlights the coherence of gene expression programs by revealing groups of DEGs that are co-regulated or co-expressed, supporting their functional relatedness and reinforcing their role as coordinated markers of the cluster.

#### 2i Validation: Classifier model with independent dataset

A Random Forest classifier is trained using the top differentially expressed genes to predict cell type labels in an independent scRNA-seq dataset containing similar populations. This approach tests whether the identified DEGs generalize across datasets and retain their discriminative power beyond the original data, providing validation that the markers capture consistent and transferable biological signals.

#### 2j Validation: Biological relevance via gene ontology enrichment analysis

To evaluate the biological functions represented in the DEGs, the ‘g:Profiler’ Python API (Reimand et al., 2007) is used to perform GO enrichment analysis. This helps link clusters to known biological processes, offering interpretability and further validating the biological relevance of the transcriptional programs observed.

### 3 Co-expression Network Analysis

To perform CNA, Weighted Gene Co-expression Network Analysis (WGCNA) is applied using the hdWGCNA R package (Morabito et al., 2023). This package is specifically adapted for single-cell data and works well with Seurat objects (Butler et al., 2018).

#### 3a Constructing metacells

Single-cell RNA sequencing data are inherently sparse due to the stochastic nature of mRNA capture and sequencing dropouts, where many genes are recorded as zero even if they are expressed at low levels. This sparsity can obscure biological signals and hinder correlation- or network-based analyses that rely on stable gene-expression measurements. To mitigate this issue, metacells are constructed; aggregate representations of transcriptionally similar cells that reduce noise while preserving biological structure.

In this study, metacells are generated using the ‘MetacellsByGroups’ procedure.

This approach first partitions the dataset into groups of similar cells using the k-Nearest Neighbors (KNN) algorithm. For each cell *i*, the algorithm identifies the set of its *k*nearest neighbors, denoted as *N*_*k*_(*i*), based on pairwise similarity in the reduced-dimensional (e.g., PCA) space. Cells that share mutual neighborhoods or belong to the same cluster are then aggregated to form a single metacell.

The expression profile of each metacell *m* is computed by averaging or summing the normalized expression values of all cells *i* that belong to it.

If 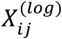 represents the log-transformed expression of gene *j* in cell *i*, and *C*_*m*_ is the set of cells assigned to metacell *m* with ∣ *C*_*m*_ ∣ members, then the metacell-level expression is given by:

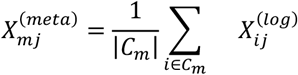

where:

- 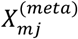 = mean expression of gene *j* in metacell *m*
- ∣ *C*_*m*_ ∣ = number of single cells aggregated into metacell *m*

Alternatively, summing rather than averaging can be used when constructing pseudobulk-like representations, depending on the requirements of downstream analyses such as co-expression network construction.

#### 3b Selecting soft-power threshold

To infer gene co-expression relationships, hdWGCNA constructs a gene–gene correlation matrix, which quantifies pairwise relationships between all genes based on their expression patterns across metacells. For each pair of genes *i* and *j*, the Pearson correlation coefficient is calculated to measure the linear association between their expression vectors. However, not all correlations represent meaningful biological connections; weak correlations often arise from noise or random fluctuations. To emphasize strong relationships and suppress weak ones, the correlation matrix is transformed into an adjacency matrix using a *soft-thresholding* procedure.

In hdWGCNA, this transformation is achieved by raising the absolute value of each correlation to a user-defined power, known as the soft-power threshold (*β*):

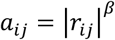

where:

- *r*_*ij*_ = Pearson correlation between genes *i* and *j*
- *a*_*ij*_ = adjacency value representing connection strength between *i* and *j*
- *β* = soft-power threshold (typically between 4 and 12)

#### 3b Constructing co-expression network

Once the soft-power threshold has been determined, hdWGCNA constructs a weighted **gene co-**expression network to identify groups of genes that exhibit coordinated expression patterns across metacells. This step captures the underlying transcriptional architecture of the dataset and provides the foundation for subsequent module detection and biological interpretation.

The first step involves computing pairwise gene–gene correlations across the metacell expression matrix. For each pair of genes *i* and *j*, a correlation coefficient *r*_*ij*_ is calculated using one of several possible metrics, Pearson, Spearman, or midweight bicorrelation, depending on the data distribution and desired robustness to outliers.

Pearson correlation is most commonly used and is defined as:

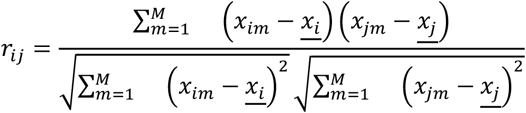

where:

- *x*_*im*_ and *x*_*jm*_ = expression levels of genes *i* and *j* in metacell *m*
- 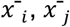 = mean expression of each gene across all *M* metacells
- *r*_*ij*_ ∈ [−1,1] = strength and direction of linear co-expression between the two genes These correlations form the similarity matrix, which is then converted into an adjacency matrix by raising each correlation’s absolute value to the previously selected soft-power threshold *β*:

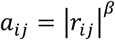

The adjacency matrix *A* = [*a*_*ij*_] represents the connection strength between genes and serves as the weighted foundation of the co-expression network. However, simple pairwise correlations may not fully capture the shared neighborhood structure between genes.

To enhance biological interpretability and network robustness, hdWGCNA transforms the adjacency matrix into a Topological Overlap Matrix (TOM). The TOM measures the similarity between the connectivity profiles of two genes; that is, how many neighbors they share within the network. Two genes that are not directly correlated but share many mutual connections will therefore have high topological overlap, indicating potential functional relatedness.

The TOM transformation emphasizes **modules of tightly connected genes** while suppressing random pairwise correlations, resulting in a network that better reflects the modular organization of transcriptional regulation.

This process is executed via the ‘ConstructNetwork’ function in hdWGCNA, which internally calls WGCNA’s core function ‘blockwiseConsensusModules’ (Langfelder & Horvath, 2008).

This implementation allows the network to be built in blocks, enabling efficient processing of large gene sets while maintaining consistent network topology across modules.

#### 3d Detecting modules

The next step in hdWGCNA is module detection; the identification of gene groups that exhibit similar expression patterns across metacells. Each module represents a cluster of genes that are highly interconnected in the co-expression network and likely participate in shared biological processes or regulatory pathways.

The TOM serves as a similarity matrix between all gene pairs, where higher values indicate greater topological overlap. To group genes with similar connectivity profiles, hdWGCNA converts this similarity matrix into a dissimilarity measure suitable for hierarchical clustering.

The TOM-based dissimilarity is defined as:

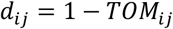

where:

- *T0M*_*ij*_ = topological overlap between genes *i* and *j*
- *d*_*ij*_ = pairwise dissimilarity, with smaller values indicating stronger co-expression relationships

Using the dissimilarity matrix *D* = [*d*_*ij*_], hierarchical clustering is performed via average linkage clustering, which iteratively merges genes or clusters based on the average pairwise dissimilarity between their members. This produces a hierarchical dendrogram where genes with similar expression connectivity profiles cluster together along tree branches.

To delineate discrete, biologically meaningful modules from the hierarchical tree, hdWGCNA employs dynamic tree cutting, an adaptive algorithm that identifies cluster boundaries by analyzing branch structure and local dissimilarity patterns.

Unlike fixed-height cutoffs, dynamic tree cutting adjusts automatically to the shape and depth of the dendrogram, allowing it to detect both large, broad modules and smaller, tightly connected gene groups within the same dataset. Each resulting module is assigned a unique color label (e.g., “turquoise,” “blue,” “brown,” etc.), consistent with the WGCNA framework. Genes not strongly associated with any module are typically grouped into a “gray” category, indicating that they did not cluster clearly with any module.

#### 3e Computing module eigengenes (ME)

Once modules have been identified, their collective gene expression profiles can be summarized by a single representative feature known as the Module Eigengene. The ME captures the dominant expression pattern of all genes within a module and serves as a lower-dimensional proxy for module activity across samples or cells. This concept, central to the WGCNA framework, allows downstream analyses to focus on the relationships between modules rather than on individual genes, thereby simplifying the network structure while retaining biological interpretability. To compute module eigengenes, Principal Component Analysis is performed on the subset of the normalized expression matrix corresponding to each gene module.

Let 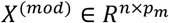 represent the expression matrix for module *m*, where *n* is the number of metacells (or single cells) and *p*_*m*_ is the number of genes in the module. PCA decomposes this matrix into principal components that capture orthogonal axes of variation among the genes:

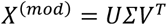

where:

- 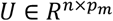 = left singular vectors (cell embeddings)
- *Σ* = diagonal matrix of singular values (variance explained)
- *V*^*T*^ = right singular vectors (gene loadings)

The first principal component (PC_1_) corresponds to the direction of maximal variance across all genes in the module and is taken as the Module Eigengene (ME):

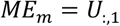

where *U*_:,1_ denotes the first column of *U*, representing the projection of each metacell (or cell) onto PC_1_ for module *m*. This vector effectively summarizes the overall expression trend of the module’s genes and serves as a quantitative measure of module activity.

hdWGCNA implements this process through the ‘ModuleEigengenes’ function, which computes MEs for each module across all cells or metacells. Additionally, Harmony batch correction can be applied directly to the eigengenes to produce harmonized Module Eigengenes (hMEs). This optional correction removes batch-related variation from eigengene values while preserving biologically relevant module-level signals, ensuring that module comparisons across conditions or datasets are not confounded by technical artifacts.

#### 3f Computing module connectivity (kME)

In co-expression network analysis, not all genes within a module contribute equally to its overall expression pattern. Some genes are more central, exhibiting strong co-expression with many other genes in the module and with the module’s eigengene itself. These highly connected and representative genes are referred to as hub genes. Hub genes are of particular biological interest because they often act as regulatory or structural anchors within molecular pathways and can provide mechanistic insight into module function.

To quantify the connectivity of each gene within a module, hdWGCNA computes the module eigengene-based connectivity, denoted as *kME*. This metric measures the correlation between the expression profile of each gene and the module eigengene that summarizes the expression of the entire module.

For a given gene *i* and module *m*, the module connectivity is defined as:

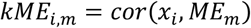

where:

- *x*_*i*_ = expression vector of gene *i*across all cells or metacells
- *ME*_*m*_ = eigengene representing module *m*
- *kME*_*i,m*_ = correlation coefficient between gene *i*and module *m*

The value of *kME*_*i,m*_ ranges from –1 to 1, with higher absolute values indicating stronger co-expression between the gene and the module eigengene. Genes with large positive *kME* values are strongly associated with the core expression pattern of the module and are typically considered intramodular hub genes.

hdWGCNA computes *kME* values using the ‘ModuleConnectivity’ function, which calculates the correlation between all genes and module eigengenes across the full single-cell dataset rather than the metacell-aggregated data. This ensures that hub gene inference captures fine-grained variation at the single-cell level while leveraging the stability of module definitions derived from metacells.

#### 3g Visualization: Dendrogram

Dendrograms are a fundamental visualization tool for representing **hierarchical clustering** results in gene co-expression network analysis. In the context of hdWGCNA, the dendrogram provides an intuitive, tree-like depiction of how genes group together based on their pairwise topological similarity derived from the Topological Overlap Matrix. This visualization helps to reveal the hierarchical structure of gene relationships; how individual genes merge into modules, and how modules relate to each other. Each leaf node on the dendrogram corresponds to a single gene, while **branches** represent clusters of genes that are progressively merged during the hierarchical clustering process.

The height on the y-axis indicates the **dissimilarity** between clusters being joined, typically computed as:

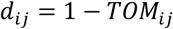

where:

- *T0M*_*ij*_ = topological overlap between genes *i* and *j*
- *d*_*ij*_ = pairwise dissimilarity used for clustering

Genes that are more similar (i.e., have higher TOM values and lower dissimilarity) are merged lower in the tree, forming tightly connected subclusters. Conversely, branches that merge higher up in the dendrogram represent gene groups that are more distinct from one another. The dendrogram is generated as part of the module detection process (Section 3d) using hierarchical clustering, where genes are iteratively grouped based on their pairwise dissimilarity. The resulting structure provides a visual summary of how genes coalesce into modules of coordinated expression. Color bands or labels are often displayed beneath the dendrogram to denote module assignments (e.g., “turquoise,” “blue,” “brown,” etc.), allowing easy identification of the boundaries between modules.

#### 3h Visualization: Module graph

Module graphs are plotted to represent each gene as a node and correlation as edge, which shows the interconnectivity of the module.

#### 3i Visualization: Module eigengene expression on UMAP

Module eigengene values are projected onto the UMAP embedding to visualize spatial or cluster-specific module activity. This allows for intuitive interpretation of module function across different cell populations or spatial contexts.

#### 3j Visualization: Module eigengene expression dot plot

Dot plots of mean module eigengene expression across cell clusters are generated to highlight patterns of activation and association with specific cell types or regions.

#### 3k Visualization: Module eigengene expression track plot

Track plots of module eigengene expression across individual cells and clusters are generated to show heterogeneity of module expression patterns.

#### 3l Visualization: Heatmap of Jaccard Index

To compare the overlap between genes identified through differential expression analysis and those belonging to co-expression network modules, a heatmap of pairwise Jaccard indices is generated. This visualization provides a quantitative and intuitive summary of how the results of DEA focused on identifying individual marker genes distinguishing clusters relate to the broader module structures identified through co-expression analysis. In other words, it bridges the gap between gene-level and network-level representations of biological organization.

For a given cluster *c* and a co-expression module *m*, let *G*_*c*_ denote the set of DEGs (differentially expressed genes) identified for that cluster, and *G*_*m*_ the set of genes belonging to module *m*.

The Jaccard Index between the two sets is defined as:

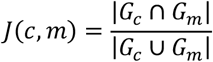

where:

- ∣ *G*_*c*_ ∩ *G*_*m*_ ∣ = number of genes shared between the cluster’s DEGs and the module
- ∣ *G*_*c*_ ∪ *G*_*m*_ ∣ = total number of unique genes present in either set
- *J*(*c, m*) ∈ [0,1], with *J* = 1 representing complete overlap and *J* = 0 indicating no shared genes

The resulting Jaccard similarity matrix *J* = [*J*(*c, m*)] quantifies the degree of correspondence between DEA-derived clusters and WGCNA-derived modules.

This matrix is visualized as a heatmap, where:

- Rows represent clusters (from differential expression analysis).
- Columns represent modules (from co-expression analysis).
- Each cell’s color intensity corresponds to the Jaccard Index value *J*(*c, m*), reflecting the proportion of shared genes.

While the Jaccard Index provides an intuitive, normalized measure of overlap between DEG lists, it does not account for background prevalence or total gene universe size. Alternative approaches such as the Fisher’s exact (hypergeometric) test incorporate these factors and yield significance values, but can be biased by highly unbalanced set sizes. In this study, the Jaccard Index was chosen as a descriptive similarity metric to visualize shared gene signatures across clusters and modules, providing an interpretable measure of proportional overlap rather than a statistical enrichment test.

#### 3m Visualization: Heatmap of connectivity

The connectivity (kME) of each gene with a module is calculated by the correlation between the gene and the module eigengene. Heatmap of kME shows how the highly connected genes correlate with different modules.

#### 3n Specificity: Tau score

To assess how selectively each module eigengene is expressed across clusters, the Tau score is computed at the module level, analogous to how it is used for individual differentially expressed genes (DEGs). While gene-level Tau scores quantify how specific a single gene’s expression is to one cluster, module-level Tau scores measure the specificity of entire co-expression programs, indicating whether a module’s activity is confined to a single cell type or broadly distributed across multiple clusters.

For each module eigengene *ME*_*m*_, let *x*_*i*_ represent its average expression (eigengene value) within cluster *i, x*_*max*_ denote the maximum eigengene value across all clusters, and *n* be the total number of clusters. The Tau score is defined as:

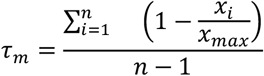

where:

- *τ*_*m*_ = Tau specificity score for module *m*
- *x*_*i*_ = mean eigengene expression of module *m* in cluster *i*
- *x*_*max*_ = *max* (*x*_1_, *x*_2_, …, *x*_*n*_) = highest eigengene expression across clusters
- *n* = number of clusters

The Tau score ranges between 0 and 1:

- *τ*_*m*_ = 0: the module eigengene is ubiquitously expressed, showing little cluster specificity.
- *τ*_*m*_ = 1: the module eigengene is perfectly specific, expressed predominantly in one cluster and absent from others.
- Intermediate values indicate partial specificity, reflecting modules active in a limited subset of clusters.

#### 3o Validation: Statistical robustness via module preservation

To evaluate the stability and reproducibility of gene co-expression modules identified by hdWGCNA, a module preservation analysis is performed. This validation step assesses whether modules discovered in one dataset (the *reference*) are preserved in another independent or resampled dataset (the *test*), thereby determining whether the modules represent stable and biologically meaningful transcriptional programs rather than artifacts of sampling or noise. The analysis follows a subsampling-based strategy. First, the co-expression network is constructed on the full dataset using the previously selected parameters (e.g., soft-power threshold, minimum module size, and tree-cutting method) to define modules at maximum resolution. These modules serve as the *reference network*. Next, the same network construction process is repeated on an independent dataset; either a distinct biological replicate or a randomly subsampled subset of the original data to create a *test network*. The goal is to compare how well the topological structure and gene–gene relationships from the reference modules are preserved in the test dataset.

The function ‘hdWGCNA::modulePreservation’ implements this analysis by quantifying multiple preservation statistics introduced in the WGCNA framework (Langfelder & Horvath, 2008). Among these, the two most informative measures are the Z-summary score and the median rank.

#### Z-summary score

Combines several preservation statistics (e.g., density and connectivity preservation) into a single composite measure that reflects the overall degree of module reproducibility.

It is defined conceptually as:

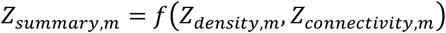

where:

- *Z*_*density,m*_ measures how densely connected the genes in module *m* remain in the test network.
- *Z*_*connectivity,m*_ measures whether the pattern of intramodular connections is preserved.
- *Z*_*summary,m*_ aggregates these into a combined preservation statistic for module *m*.

Interpretation of the Z-summary score follows the standard WGCNA convention:

- *Z*_*summary,m*_ > 10: strongly preserved module
- 2 < *Z*_*summary,m*_ ≤ 10: moderately preserved module
- *Z*_*summary,m*_ ≤ 2: not preserved

#### Median rank

Ranks modules according to their preservation metrics, with lower ranks indicating stronger preservation. This measure is less dependent on module size and provides a relative comparison among modules within the same dataset. Modules with high Z-summary scores and low median ranks are interpreted as robust, meaning that their co-expression structure remains consistent when evaluated on independent data. Such preservation indicates that these modules capture reproducible transcriptional programs, rather than dataset-specific or noise-driven patterns. In practice, hdWGCNA’s ‘modulePreservation’ function performs permutation testing to estimate empirical null distributions for preservation statistics, thereby providing significance estimates for each metric. This rigorous validation ensures that modules used for biological interpretation or downstream analyses represent stable, generalizable regulatory programs that can be consistently recovered across independent samples or experimental conditions.

#### 3p Validation: Biological relevance via gene ontology enrichment analysis

Beyond assessing statistical robustness, it is essential to evaluate whether the identified co-expression modules correspond to biologically meaningful pathways or cellular functions. To achieve this, each module is tested for enrichment of Gene Ontology (GO) terms and other functional annotation categories using hdWGCNA::RunEnrichR, which interfaces with the *EnrichR* platform (Chen et al., 2013). This step provides biological validation, ensuring that the modules represent coherent transcriptional programs rather than random collections of co-expressed genes.

For each module *m*, the set of its member genes *G*_*m*_ is submitted to EnrichR to identify GO terms, pathways, or curated gene sets that are statistically overrepresented relative to a background list (typically all genes tested in the network). Enrichment analysis evaluates whether the overlap between *G*_*m*_ and a given annotation category *A* exceeds what would be expected by chance. This is formally assessed using the Fisher’s exact test (or equivalently, the hypergeometric test), where the probability of observing at least *k* overlapping genes is computed as:

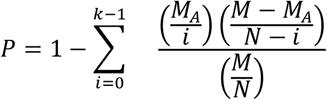

where:

- *M* = total number of genes in the background set
- *M*_*A*_ = number of genes annotated to category *A*
- *N* = number of genes in module *m*
- *k* =∣ *G*_*m*_ ∩ *A* ∣ = number of overlapping genes between the module and the annotation category

The resulting *P*-values are adjusted for multiple testing using the Benjamini–Hochberg False Discovery Rate (FDR) method to control for false positives. Categories with adjusted *P*-values below a predefined threshold (typically FDR < 0.05) are considered significantly enriched. The output of ‘RunEnrichR’ includes a ranked list of enriched biological terms for each module, along with corresponding significance values, enrichment scores, and gene overlap counts. Visualization of enrichment results, such as bar plots or bubble plots, highlights the top biological processes, molecular functions, or cellular components associated with each co-expression module.

#### 3.5.2 Data selection

The Allen Brain Cell Atlas dataset used in this study was obtained from the Allen Institute for Brain Science, which provides a comprehensive, hierarchically annotated taxonomy of mouse brain cell types (Yao et al., 2023). The ABCA taxonomy was derived from approximately 1.3 million cells sampled across multiple cortical and hippocampal regions and profiled using two independent sequencing platforms (10x Genomics and SMART-Seq). Cell-type annotations were generated through an iterative process of unsupervised clustering, marker gene identification, and expert curation, yielding a multi-level hierarchy that includes class, subclass, and supertype designations. To ensure the robustness of these labels, the ABCA integrated cross-platform validation, demonstrating strong agreement between 10x and SMART-Seq datasets, and performed orthogonal spatial validation using spatial transcriptomics to confirm the anatomical localization of transcriptomic clusters within known brain structures. These cross-modal and cross-platform validations provide high confidence in the reproducibility and biological accuracy of the ABCA annotations. While all reference atlases have inherent limitations in scope and coverage, the ABCA remains one of the most rigorously validated resources available for defining mouse brain cell types and serves as a trusted benchmark for evaluating computational frameworks of cellular identity in this study.

ABCA comprises large-scale, single-cell-resolution transcriptomic datasets spanning all brain regions, generated using single-cell RNA sequencing (Yao et al., 2023). The scRNA-seq datasets were generated using 10x Genomics Chromium versions 2 and 3 (10xv2 and 10xv3) from anatomically defined tissue microdissections, resulting in approximately 7.0 million single-cell transcriptomes, representing around 5% of all cells in the mouse brain. After applying stringent quality control (QC) criteria, low-quality transcriptomes were excluded, leaving 4.3 million QC-passed cells. These cells underwent iterative clustering, resulting in an initial transcriptomic taxonomy comprising 5,283 clusters. To identify discriminative features among clusters, all pairwise comparisons were performed within this taxonomy, yielding 8,460 differentially expressed genes (DEGs). To hierarchically organize the taxonomy and define relationships among clusters, Pearson correlation coefficients of gene expression were computed between all cluster pairs using either all DEGs or subsets thereof. These analyses revealed varying degrees of similarity among clusters, which could be grouped into broader categories. Notably, transcription factor marker genes yielded the lowest correlation values across the brain compared to functional markers, adhesion molecules, or all marker genes, and therefore provided the best resolution of global cluster relationships. Based on these results, the transcription factor marker genes were used to computationally construct a hierarchical organization of the taxonomy, classifying clusters into putative classes, subclasses, and supertypes.

The ABCA provides a comprehensive platform for visualizing and analyzing multimodal single-cell data across the mammalian brain. Data associated with the ABCA is hosted on Amazon Web Services (AWS) in an S3 bucket as a public dataset (arn:aws:s3:::allen-brain-cell-atlas), accessible without the need for an account or login. To facilitate data retrieval and management, we employed the AbcProjectCache, a lightweight Python object provided by the Allen Institute. This tool streamlines the downloading process and manages different data release versions. Detailed documentation and example use cases for the AbcProjectCache are available in the Allen Brain Cell Atlas Data Access repository (https://alleninstitute.github.io/abc_atlas_access/intro.html).

## Notes

Conflicts of interest: The authors have no competing financial interests.

### Competing Interest Statement

The authors have declared no competing interest.

